# Illuminating stomatal responses to red light: establishing the role of *C_i_*-dependent vs - independent mechanisms

**DOI:** 10.1101/2023.10.27.564341

**Authors:** Georgia Taylor, Julia Walter, Johannes Kromdijk

**Affiliations:** Department of Plant Sciences, University of Cambridge, Cambridge, Cambridgeshire, CB23EA, UK; Carl R Woese Institute for Genomic Biology, University of Illinois at Urbana-Champaign, IL61801, Illinois, USA

**Keywords:** stomatal conductance, guard cells, photosynthesis, intercellular CO2 (*C_i_*), red-light, signalling, stomata

## Abstract

The stomatal response to red light appears to link stomatal conductance (*g_s_*) with photosynthetic rates. Initially, it was suggested that changes in intercellular CO *_2_* (*C_i_*) provide the main cue via a *Ci-* dependent response. However, evidence for *C_i_*-independent mechanisms suggests an additional, more direct relationship with photosynthesis. While both *C_i_*-dependent and -independent mechanisms clearly function in stomatal red-light responses, little is known regarding their relative contribution. The present study aimed to quantify the relative magnitude of *C_i_*-dependent and - independent mechanisms on the stomatal red-light response, to characterise their interplay and to assess the putative link between plastoquinone (PQ) redox state and *C_i_*-independent stomatal responses. Red light response curves measured at a range of *C_i_* values for wild-type *Arabidopsis thaliana* (Col-0) and the CO*_2_* hyposensitive mutant, *ca1ca4*, allowed deconvolution of *C_i_*-dependent and -independent pathways. Surprisingly, we observed that both mechanisms contribute equally to stomatal red-light responses, but *C_i_*-independent stomatal opening is suppressed at high *C_i_*. The present data are also consistent with the involvement of PQ redox in coordinating *C_i_-*independent stomatal movements. Overall, it seems that while *C_i_*-independent mechanisms are distinct from responses to *C_i_,* interplay between these two pathways is important to facilitate effective coordination between *g_s_* and photosynthesis.

**Highlight:** By controlling intercellular CO_2_ (*C_i_*) at a range of values across red-light response curves, we demonstrate independent and interactive roles of *C_i_-*dependent and –independent pathways in coordinating stomatal red-light responses.

## Introduction

Stomata are microscopic pores on the leaf epidermis which govern the pathway of carbon dioxide and water vapour exchange between plants and the surrounding atmosphere. Highly specialised guard cells (GCs) surround each pore and respond to various environmental and endogenous cues, such as light, CO*_2_*, humidity and abscisic acid (ABA), to regulate stomatal conductance (*g_s_*) via changes in GC turgor pressure (Roelfsema *et al*., 2002; Inoue & Kinoshita, 2017; Jezek & Blatt, 2017; Lawson & Matthews, 2020). As such, GC responses are crucial for balancing photosynthetic demand for carbon with the need to conserve water (Buckley, 2017; Wong *et al.,* 1979). However, some of the exact mechanisms by which *g_s_* and net CO*_2_* assimilation (*A_net_*) are coordinated remain unclear.

Since light intensity and spectral quality are important determinants of *A_net_*, effective coordination between *g_s_* and *A_net_* requires *g_s_* to respond to the prevailing light conditions. Stomatal responses to light can be divided into two distinct wavelength-dependent pathways. Blue light is perceived directly at the guard cells via two blue light photoreceptor proteins called phototropins (*PHOT1* and *PHOT2*), inducing a well-characterised signal transduction cascade that leads to rapid membrane hyperpolarisation and stomatal opening even at low fluences (Shimazaki *et al*., 1986; Kinoshita *et al.,* 2001; Shimazaki *et al.,* 2007; Takemiya *et al.,* 2013; Inoue & Kinoshita, 2017). By contrast, much less is known about the mechanisms which underpin stomatal responses to red light. Also known as the ‘quantitative’ light response, the red-light response is slower than the response to blue light and occurs at higher irradiances, saturating at similar light intensities to photosynthesis (Sharkey & Raschke, 1981). Red light-induced stomatal opening is abolished by inhibitors of photosynthetic electron transport, such as 3-(3,4-dichlorophenyl)-1,1-dimethylurea (DCMU) (Sharkey & Raschke, 1981; Olsen *et al.,* 2002; Messinger *et al.,* 2006; Wang *et al.,* 2011; Ando & Kinoshita, 2018), which suggests red-light stomatal responses are directly impacted by signals derived from photosynthesis (Lawson *et al.,* 2008; Matthews *et al.,* 2020). However, the exact location and mechanisms by which these putative photosynthesis-related signals arise and are communicated to inflict changes in GC turgor remain elusive.

Early work suggested that stomatal responses to intercellular CO *_2_* (*C_i_*) provide the main cue for red light-induced stomatal opening, indirectly coupling *g_s_* with the mesophyll’s demand for CO *_2_*(Roelfsema *et al.,* 2002, 2006). Recent genetic advances have provided insight into the molecular mechanisms underlying this so-called ‘ *C_i_-*dependent response’. The *C_i_*-dependent response can be divided into two CO*_2_*-concentration-dependent pathways, in which stomatal closure is promoted by elevated *C_i_* concentrations, whereas reductions in *C_i_* provide a potent cue for stomatal opening (Mott, 1988; Fujita *et al.,* 2013; Engineer *et al.,* 2016; Dubeaux *et al.,* 2021). Within these opposing *C_i_*-dependent responses, multiple pathways exist and converge to modulate GC behaviour. Low *C_i_*concentrations promote stomatal opening via the RAF-like MAP kinase kinase kinase, HIGH LEAF TEMPERATURE1 (HT1), which negatively regulates stomatal closure (Hashimoto *et al.,* 2006; Matrosova *et al.,* 2015; Hashimoto-Sugimoto *et al.,* 2016). Although the exact signalling cascade downstream of HT1 is not fully elucidated, two RAF-like kinases, CONVERGENCE OF BLUE LIGHT (CBC) 1 and 2, appear to be involved (Matrosova *et al.,* 2015; Hiyama *et al.,* 2017). Active HT1 has been shown to interact with and phosphorylate CBC1/2 *in vitro* (Hiyama *et al.,* 2017) and it is proposed that following phosphorylation, active CBC1/2 inhibit guard cell S-type anion channels, such as SLAC1, to suppress stomatal closure (Matrosova *et al.,* 2015; Hiyama *et al.,* 2017). Low CO*_2_*also promotes the phosphorylation and activation of plasma membrane (PM) H *^+^*-ATPase channels (Kinoshita & Shimazaki, 1999, 2002; Kinoshita *et al*., 2001), which in turn promote stomatal opening via activation of voltage-gated inward-rectifying K *^+^* channels (Schroeder *et al.,* 1987).

By contrast, high *C_i_* concentrations are perceived within the guard cells via two beta carbonic anhydrases, CARBONIC ANHYDRASE 1 (CA1) and CA4, which strongly accelerate the equilibration between CO*_2_* and bicarbonate (HCO*_3_^-^*) (Hu *et al.,* 2010; Xue *et al.,* 2011). In *Arabidopsis,* high CO*_2_*-induced stomatal closure is suppressed in plants lacking both enzymes (Hu *et al.,* 2010) and could be restored by complementation with either CA1, CA4 or an unrelated mammalian αCAII, suggesting that the catalytic activity of these CAs is required for high-CO *_2_* induced stomatal closure (Hu *et al.,* 2010). In addition, the MAP kinases MPK4 and MPK12 were recently identified as downstream CO *_2_*/ HCO*_3_^-^* sensors (Tõldsepp *et al.,* 2018; Takahashi *et al.,* 2022; Yeh *et al.,*2023). In response to elevated bicarbonate MPK4/12 directly interact with and inhibit HT1, which in turn relieves HT1-mediated repression of GC S-type anion channels and promotes stomatal closure (Takahashi *et al.,* 2022; Yeh *et al.,* 2023). In addition to the HT1-mediated pathway of stomatal closure, high CO*_2_* also promotes rapid dephosphorylation and deactivation of PM H *^+^*-ATPase channels, which suppresses stomatal opening (Edwards & Bowling, 1985; Ando *et al.,* 2022).

While changes in *C_i_* provide a potent cue for stomatal opening and closing, an increasing body of evidence suggests that the stomatal red-light response may also be controlled by *C_i_*-independent mechanisms. In particular, stomata continue to respond to changes in red light intensity when *C_i_* is kept constant, suggesting alternative red light sensing and signalling pathways must also be involved (Messinger *et al.,* 2006; Lawson *et al.,* 2008; Wang & Song, 2008). The maintenance of red light-induced stomatal opening in the CO *_2_* hyposensitive mutant, *ca1ca4*, provides further evidence for a *C_i_*-independent mechanism (Matrosova *et al.,* 2015; Ando *et al.,* 2018). Based on a synthesis of experimental work, Busch (2014) suggested that the redox state of the chloroplast plastoquinone (PQ) pool may provide an early photosynthesis-derived signal to coordinate stomatal responses to red light. This putative relationship appears consistent with altered stomatal responses observed in *Nicotiana tabacum* plants with modified levels of *Photosystem II Subunit S*(*PsbS*; Głowacka *et al*., 2018). Additionally, incorporation of the hypothesised PQ redox signal improved performance of models for stomatal conductance (Kromdijk *et al*., 2019). However, the mechanisms by which these redox signals could be communicated away from the chloroplast and ultimately coordinate stomatal behaviour are not yet understood.

While there is clear evidence for the involvement of both *C_i_-*dependent and -independent mechanisms in coordinating the stomatal response to red light, several open questions remain. For example, how much does each mechanism (i.e., *C_i_-*dependent and -independent) contribute to red-light-induced stomatal opening? And are these pathways distinct or do they show interaction with one another?

In the present work, the objectives were to quantify the relative magnitude of *C_i_*-dependent and - independent mechanisms in coordinating the stomatal red-light response, to characterise their interplay across a wide range of light intensities and *C_i_* concentrations and to assess the putative involvement of PQ redox state in *C_i_*-independent responses. To decouple stomatal control via *C_i_*-dependent and -independent pathways, red light response curves were measured at a range of *C_i_* values on wild-type *Arabidopsis thaliana* (Col-0) and on the CO*_2_* hyposensitive mutant, *ca1ca4*, which allowed further deconvolution of these pathways. Somewhat surprisingly, the results indicate that the contribution of *C_i_* -dependent and -independent mechanisms to red-light-induced stomatal movements is of similar magnitude. In addition, the *C_i_*-independent opening response was found to be distinct from and additive to the *C_i_*-dependent response to red light, while the *C_i_*-dependent closing response is dominant and strongly suppresses *C_i_*-independent opening. Overall, these results provide further insight into the mechanisms which control stomatal responses to red light and provide new avenues for future work to fully elucidate the molecular regulation involved.

## Materials and methods

### Plant materials and propagation

Seeds of *Arabidopsis thaliana*carbonic anhydrase knockout mutant *ca1ca4* (Hu *et al.,* 2010) were obtained from the Nottingham Arabidopsis Stock Centre (NASC) and genotyped via PCR (for primers see Table S1). Col-0 and *ca1ca4* seeds were sown on a 4:1 mix of Levington® Advance F2 compost : sand and stratified at 4*°*C for 4 days. Seedlings were then transferred into individual 7×7 cm pots and positioned into a controlled growth chamber with a short-day photoperiod (8 h light / 16 h dark). Light intensity was controlled at ∼200 µmol m *^-1^* s*^-2^*, relative humidity (RH) at 60% and air temperature at 20*°*C. Plants were hand-watered and randomly repositioned on the growth shelf every 3-4 days. All gas exchange measurements were performed on young fully expanded leaves of 7-9-week-old plants.

### Gas exchange measurements

#### Determining Ci-dependent vs -independent responses

Col-0 or *ca1ca4* plants were dark adapted for 2 hours, after which a fully expanded leaf (rosette leaf number 7-9) was clamped into the cuvette of an open gas exchange system (LI6400XT, LI-COR) with a 2 cm*^2^* integrated fluorometer head (Leaf Chamber Fluorometer, LI6400-40, LI-COR). Block temperature was controlled at 25 *°*C, reference CO*_2_* at 410 ppm and relative humidity was maintained between 50-70%. For leaves that did not completely fill the cuvette, an image of the leaf inside the gasket was taken and leaf area was calculated using ImageJ (ImageJ, U. S. National Institutes of Health, MD, USA) and adjusted in the gas exchange calculations. The reference CO *_2_* was adjusted to obtain a defined intercellular CO *_2_* (*C_i_*) concentration (75, 150, 300, 375, 450, 600, 750 µmol mol*^-1^*), and leaves were allowed to equilibrate for at least 30 minutes in the dark at these *C_i_*concentrations, until *g_s_* had stabilised. The measuring light was then switched on briefly and once *F* stabilised, chlorophyll fluorescence and gas exchange parameters were recorded. All fluorescence measurements were obtained using the multiphase flash routine (Loriaux *et al*., 2013), with the saturating flash set to 4000 µmol m *^-2^* s*^-1^* and a ramp of 30%. The minimal (Fo) and maximal (Fm) fluorescence were used to determine the maximal efficiency of whole-chain electron transport (Fv/Fm). The actinic light (100% red light, RL) was then increased stepwise: 0, 50, 100, 200, 400, 600, 800 µmol m*^-2^* s*^-1^*. For each increase in RL, the CO*_2_* concentration of the cuvette was altered via the millivolt signal to maintain a constant *C_i_* concentration across the red-light response curve. Plants were acclimated for at least 30 minutes at each light intensity and *C_i_* concentration before recording gas exchange and chlorophyll fluorescence parameters (F’, Fo’ and Fm’). Table 1 provides the definitions and equations for the fluorescence parameters used. To account for diffusional leakage of CO*_2_* from the cuvette all gas exchange data were leak-corrected as per McDermitt *et al*. (1989) with some modifications. Briefly, leakage coefficients were calculated for each genotype using the dark-measured *A_net_* values recorded for each*C_i_* concentration. Gas exchange data (*A_net_* and *C_i_*) was then corrected using these coefficients (McDermitt *et al.,* 1989).

**Table 1.** Definitions and equations of measured and calculated fluorescence parameters (Kramer et al., 2004; Maxwell & Johnson, 2000; Murchie & Lawson, 2013)

| Parameter | Definition | Equation |
| --- | --- | --- |
| $F_o$ | Minimal fluorescence (dark-adapted) | |
| $F_m$ | Maximum fluorescence (dark-adapted) | |
| $F'$ | Steady-state fluorescence | |
| $F_o'$ | Minimal fluorescence (light-adapted) | |
| $F_m'$ | Maximal fluorescence (light-adapted) | |
| $F_v/F_m$ | Maximum quantum yield of PSII (dark-adapted) | $(F_m - F_o) / F_o$ |
| $F_q'/F_m'$<br>($\Phi_{PSII}$) | Operating efficiency of PSII | $(F_m' - F') / F_m'$ |
| NPQ | Non-Photochemical Quenching | $(F_m - F_m') / F_m'$ |
| $1 - q_L$ | $Q_A$ redox state | $q_L = (1/F' - 1/F_m') / (1/F_o' - 1/F_m')$ |

#### Determining steady-state ‘physiologically relevant’ C _i_ values

Col-0 plants were dark adapted for 2 hours, after which a fully expanded leaf (rosette leaf number 7-9) was clamped into the gas exchange cuvette. Block temperature was controlled at 25 *°*C, reference CO*_2_* at 410 ppm and relative humidity was maintained between 50-70%. Actinic red light (RL; 100%) was increased in a stepwise manner across a physiologically relevant range of light intensities: 0 (Dark; D), 50 (low-light; LL), 200 (growth-light; GL) and 800 (high-light; HL) µmol m *^-2^* s*^-1^*. The intercellular CO*_2_* (*C_i_*) values obtained in the dark (D), at LL and HL were used to determine a range of physiologically relevant *C_i_*concentrations.

### Statistical analysis

Two-way repeated measures ANOVA was used to assess the significance of effects of red-light intensity, *C_i_*, and their interaction on stomatal conductance. Additionally, η *^2^* was calculated to estimate the proportion of variance associated with each effect. For each ANOVA, assumptions of normality, homogeneity of variance and sphericity were tested. For data which failed to meet these assumptions, a non-parametric Kruskal-Wallis rank sum was used, followed by Dunn’s multiple comparison test. The relationship between Q *_A_* redox and stomatal conductance was analysed using multiple linear regression and minimum models for Col-0 and *ca1ca4* were obtained via backwards stepwise elimination. All data analysis and plot generation was carried out using R 4.1.1 (R Core Team, 2021) on RStudio (Posit Team, 2022).

## Results

### C_i_-dependent and -independent mechanisms contribute equally to the stomatal red light response

To empirically quantify the relative contributions of *C_i_-*dependent and *C_i_-*independent mechanisms on stomatal responses to red light, gas exchange measurements were performed on *A. thaliana* (Col-0) plants at a matrix of 7 red light intensities and 7 *C_i_* values to obtain red-light dose response curves of *A_net_* (Figure 1A) and *g_s_* (Figure 1B) across *C_i_*-values spanning from 75 to 750 µmol mol *^-1^*. To separate *C_i_*-dependent and *C_i_*-independent responses, *C_i_* values were kept constant within each red-light response curve (Figure 1C) by carefully adjusting the CO *_2_* concentration of the cuvette after each change in light intensity, following the approach by Messinger *et al*. (2006). In this way, the response of *g_s_* to red light intensity within each response curve can be taken as a direct measure of *C_i_*-independent mechanisms, while the separation between response curves at different *C_i_* values provides an estimate of *C_i_*-dependent mechanisms.

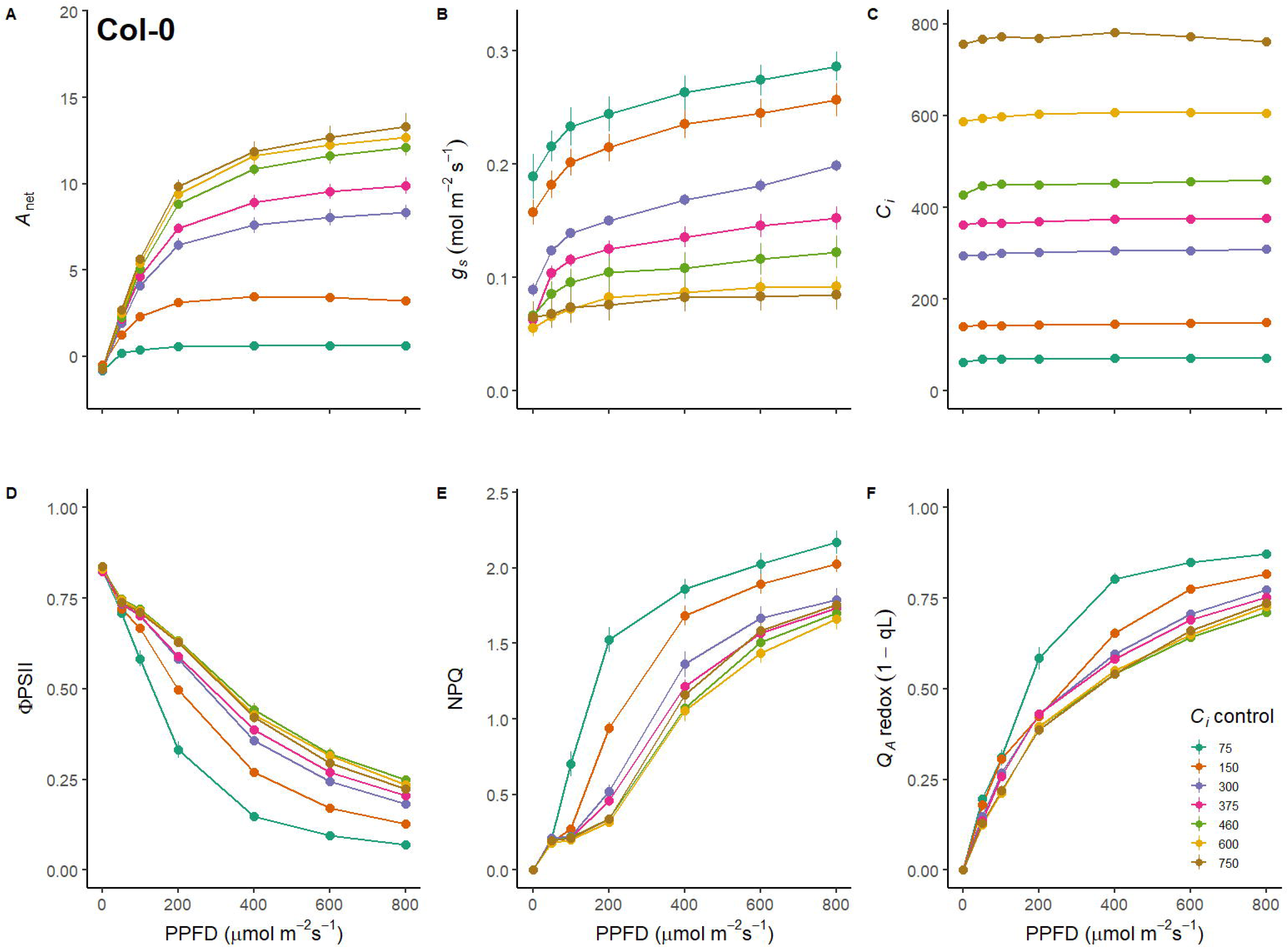

As expected, *A_net_* increased hyperbolically with increasing red-light intensity. At *C_i_* values of 75 and 150 µmol mol*^-1^*, *A_net_* was strongly inhibited by limiting CO *_2_* supply, whereas *A_net_* reached saturation for *C_i_* values above 460 µmol mol *^-1^* (Figure 1A). Interestingly, *g_s_* values were strongly influenced by red light intensity changes at each constant *C_i_* as well as by the different *C_i_* levels between the response curves (Figure 1B). Namely, while *C_i_* was kept constant across each light response curve *, g_s_* displayed a continued opening response to increasing red-light intensity at each *C_i_* level (Figure 1B). Although the general shape of this opening response remained largely consistent between the different *C_i_* values, the entire *g_s_*/light curves were transposed along the*g_s_*-axis, due to clear separation in initial (*g_s int_*) and maximal *g_s_* (*g_s max_*) resulting from the contrasting *C_i_* values. In line with these observations, two-way repeated measures ANOVA demonstrated highly significant effects of red-light intensity at constant*C_i_* (*F*(6,47) =321.93, *P* < 0.001), *C_i_* level (*F*(6,36) =39.67, *P* < 0.001) and their interaction (*F*(36,282) = 9.201, *P <* 0.001) on the response of *g_s_.* η*^2^* was calculated to estimate the proportion of variance associated with each effect. Remarkably, both light intensity (η *^2^*= 0.87) and *C_i_* (η*^2^*= 0.84) had very similar effect size, with slightly smaller effect size of the interaction (η *^2^=* 0.54).

Changes in electron transport parameters were concomitantly estimated from fluorescence measurements taken alongside gas exchange after acclimation to each light level (Figure 1D-F). The maximal efficiency of PSII after dark acclimation (*F_v_/F_m_,* Figure 1D) was 0.83 ± 0.001, similar across all measurements and was not significantly affected by *C_i_* value (*P* = 0.16). For *C_i_* values above 150 µmol mol*^-1^*, the quantum efficiency of PSII (ΦPSII) as a function of red-light intensity was similar, whereas low *C_i_* values (< 150 µmol mol*^-1^*) caused a reduction in ΦPSII (Figure 1D), consistent with carbon supply limiting CO*_2_* assimilation (Figure 1A) and consequently inhibiting linear electron flow. Likewise, carbon supply limitation also affected the NPQ response, where a greater induction of NPQ was observed at low*C_i_* values (Figure 1E). Plastoquinone redox state can be approximated by the fluorescence parameter 1 – q*_L_* (Kramer *et al*., 2004), which estimates the redox state of the first stable electron acceptor at PSII, the quinone bound to the Q *_A_* site. In line with the observed patterns of ΦPSII, Q*_A_* redox state as a function of red-light intensity was similar for *C_i_* values above 150 µmol mol*^-1^* but lower *C_i_* values caused the Q*_A_* pool to become more reduced (i.e., higher values of 1-q *_L_*, Figure 1F).

The wide range of controlled *C_i_* values used to construct the red-light response curves in Figure 1C may exceed the *C_i_* levels typically experienced by plants under steady-state non-stressed conditions. To assess the most relevant range of *C_i_* values for red-light stomatal responses in non-stressed *A. thaliana* plants, a second set of light response measurements was performed. For these measurements, the CO*_2_* concentration in the cuvette was kept at the approximate level during plant growth (410 µmol mol*^-1^*), allowing *C_i_* values to respond to each change in light intensity (Figure S1C). Concomitant with increasing *A_net_* and *g_s_* (Figure S1A & B),*C_i_* concentration declined with increasing red-light intensity (Figure S1C), ranging from 437 ± 3 µmol mol *^-1^* in darkness to 312 ± 5 µmol mol *^-1^* at a light intensity of 800 µmol m *^-2^* s*^-1^*. We therefore estimate the physiologically relevant *C_i_* range for stomatal control in these plants to be approximately between 300-450 µmol mol *^-1^*. As such, we reanalysed the relationship between stomatal conductance, *C_i_* and light intensity using only curves at *C_i_* values within this physiologically relevant range. A two-way ANOVA showed a significant effect of *C_i_* concentration (*P* < 0.002), light intensity (*P* < 0.001) and their interaction (*P* < 0.001) on stomatal conductance. Interestingly, the relative effect size of these factors shifted, whereby the contribution of light intensity remained similar (η *^2^*= 0.91), while the proportion of variance associated with *Ci* concentration (η*^2^*= 0.43) and their interaction (η*^2^*= 0.42) was considerably reduced.

### Stomatal conductance responds strongly to *C*_i_ in darkness

Interestingly, different *C_i_*-values seemed to inflict substantial changes in *g_s_* in the absence of light. Indeed, statistical analysis of *g_s int_*, which was recorded after at least 30 minutes of acclimation to the controlled *C_i_* concentration in darkness (Figure 2A), showed a significant effect o*C*f*_i_* value (Kruskal-Wallis *H* = 36.129, df = 6, *P* < 0.001). In particular, sub-ambient *C_i_* values induced a significant stomatal opening response, with approximately three-fold higher *g_s int_* at *C_i_* of 75 and 150 µmol mol*^-1^*, compared to any of the *C_i_* values greater than 375 µmol mol *^-1^* (P < 0.05, for all Dunn’s comparisons see Table S2). These results clearly demonstrate that stomatal movements via C *_i_*-dependent mechanisms occur in darkness, and that low *C_i_*concentrations are sufficient to induce significant stomatal opening independently of light.

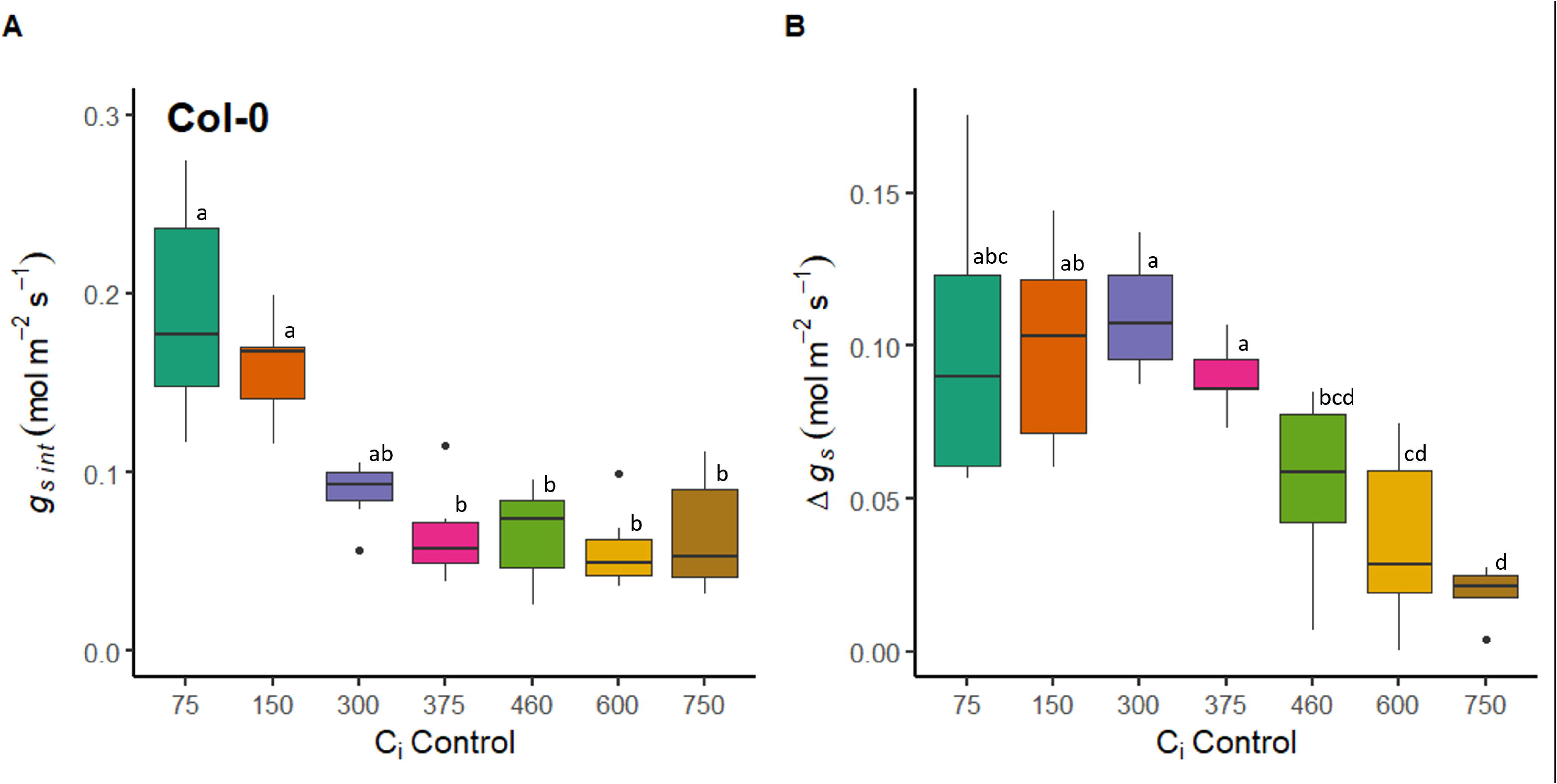

### The magnitude of *C*_i_-independent stomatal responses to red light is suppressed by high *C*_i_

To extract the response range of*C_i_*-independent responses to red light, Δ*g_s_* (= *g_s max_* – *g_s int_*) was calculated for each *C_i_* value (Figure 2B). Δ*g_s_* was significantly affected by*C_i_* value (Kruskal-Wallis *H* = 34.422, df = 6, *P <*0.001). Δ*g_s_* was approximately 0.1 mol m *^-2^* s*^-1^* and did not vary significantly for response curves measured at *C_i_* values less than 460 µmol mol *^-1^* (P < 0.05, for all Dunn’s comparisons see Table S2), demonstrating that *C_i_* values did not affect the magnitude of *C_i_*-independent mechanisms within this range. Interestingly, the most pronounced Δ *g_s_* were observed at*C_i_* values within the physiologically active range, at 300 µmol mol *^-1^*. By contrast, a reduction in Δ*g_s_* was observed in response to high *C_i_* (> 460 µmol mol*^-1^*) and the extent of Δ*g_s_* inhibition increased with *C_i_* values; such that the opening response associated with *C_i_*-independent mechanisms was almost completely abolished at a *C_i_* value of 750 µmol mol*^-1^*.

### CO_2_ hyposensitivity maintains *C*_i_-independent stomatal opening across the full *C*_i_ range

To further investigate the observed suppression of *C_i_*-independent stomatal opening at high*C_i_* concentrations in Col-0, responses of *A_net_* (Figure 3A) and *g_s_* (Figure 3B) to increasing red light under the same controlled *C_i_* treatments (Figure 3C) were measured on CO *_2_* hyposensitive *ca1ca4* plants. As expected, *A_net_* of *ca1ca4* increased with red-light intensity and was strongly inhibited at low *C_i_* and elevated above ambient at high *C_i_* concentrations (Figure 3A). Interestingly, while *ca1ca4* demonstrated a strong stomatal opening response to increasing red light (Figure 3B), the *C_i_*-dependent separation of stomatal responses was not very pronounced. Instead, stomatal responses clustered together for all *C_i_* values above 75 µmol mol *^-1^* (Figure 3B). Notably, a pronounced increase in *g_s max_* was observed at a *C_i_* value of 75 µmol mol*^-1^*, indicating that the response to low *C_i_* is at least partially maintained in this double mutant. Indeed, a two-way repeated measures ANOVA demonstrated a significant effect of *C_i_* concentration (*F*(5, 45) = 3.617, *P* = 0.005) and a highly significant effect of red-light intensity (*F*(6,36) = 369.179, *P* <0.001), and their interaction (*F*(36, 270) = 3.558, *P* <0.001) on the stomatal response. Interestingly, when the proportion of variance associated with each variable in *ca1ca4* was calculated, the contribution of light intensity remained high (η*^2^*= 0.89), while the effect sizes of *C_i_* (η*^2^*= 0.33) and the interaction (η*^2^=* 0.16) decreased considerably due to the CO*_2_* hyposensitivity in this genotype.

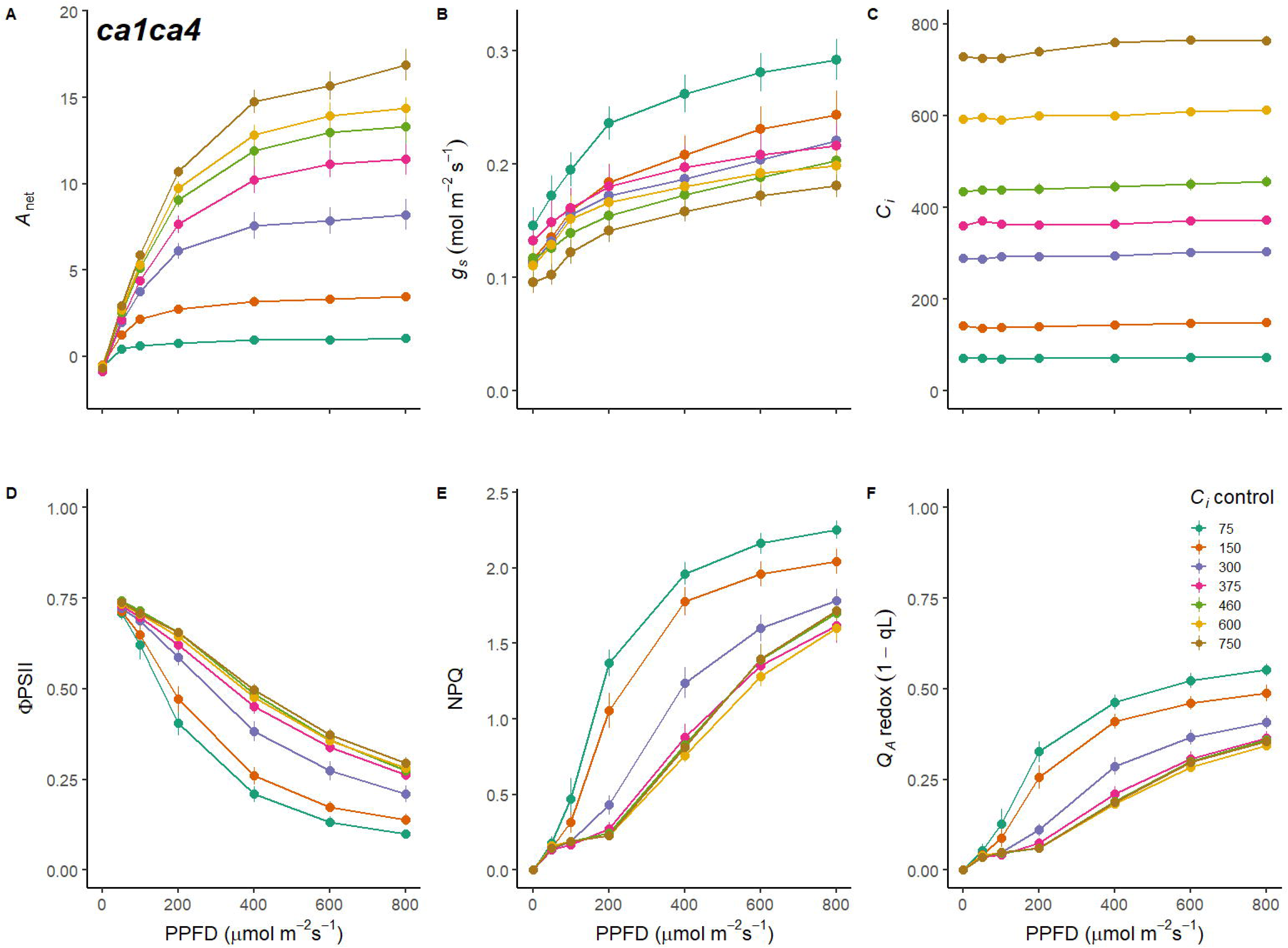

Carbon supply limitation influenced the responses oΦf PSII, NPQ and Q*_A_* redox to light intensity (Figure 3D-F). *F_v_/F_m_* was not significantly affected by *C_I_* value (*F_v_/F_m_* = 0.831 ± 0.002, *P* = 0.292). However, for *C_i_* concentrations of <300 µmol mol *^-1^* a pronounced reduction of ΦPSII was observed (Figure 3D), with a concomitant increase in NPQ activation (Figure 3E) and Q *_A_* reduction (Figure 3F).

### *C*_i_-dependent responses are impaired in *ca1ca4*

Interestingly, *C_i_-*induced stomatal movements under dark conditions were not significant in *ca1ca4* (Figure 4A; *F*(6, 45) = 1.072, *P* = 0.394), suggesting that the activity of CA1/CA4 is crucial for the sensing and coordinating of CO*_2_* -induced stomatal movements in the absence of light.

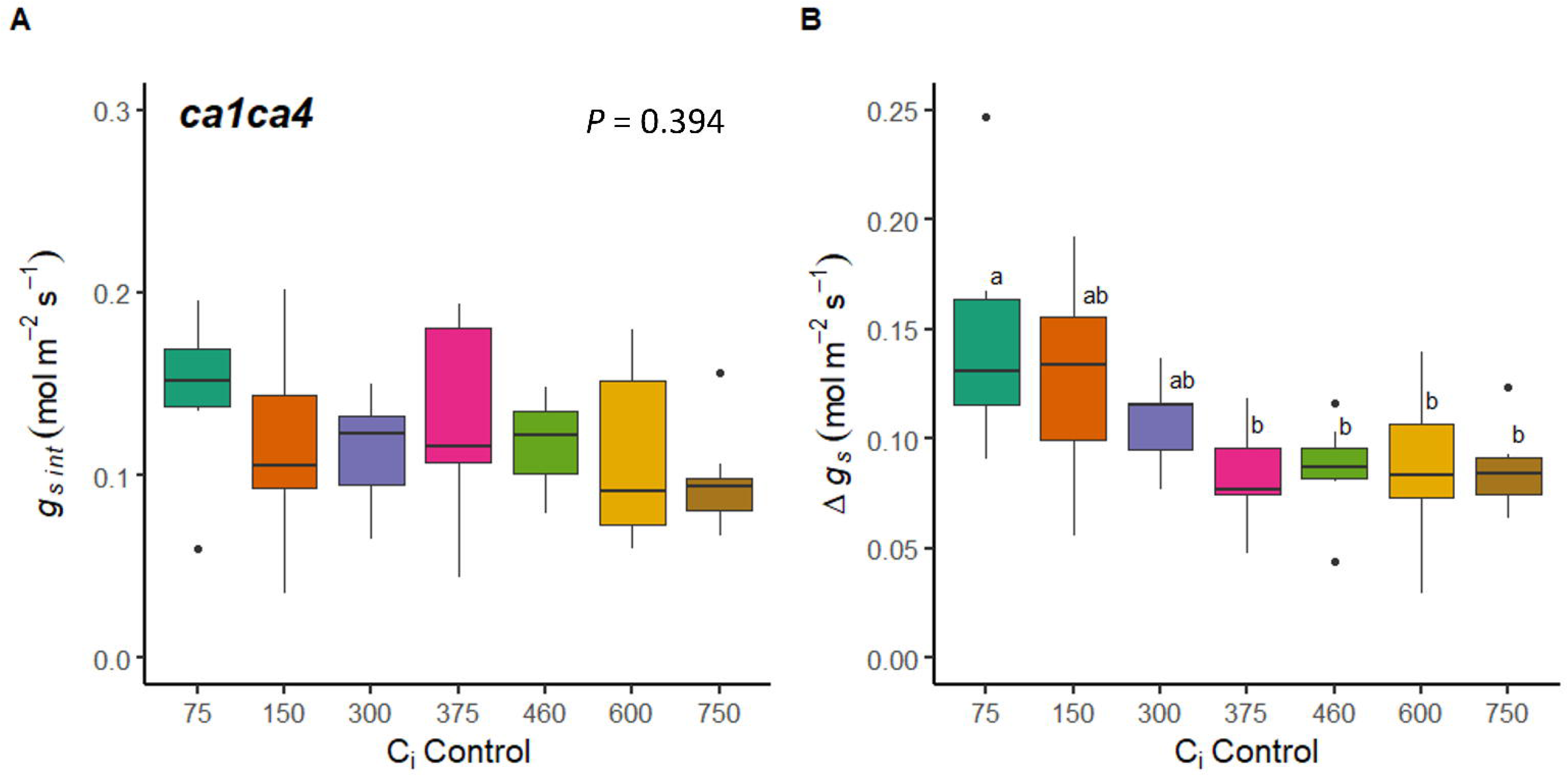

To investigate the effect of CO*_2_* hyposensitivity on the magnitude of *C_i_*-independent stomatal opening, we also compared Δ*g_s_* across all *C_i_* concentrations for*ca1ca4* (Figure 4B). While Δ*g_s_* was still significantly affected by *C_i_* concentration (*F*(6,45)=3.98, *P* = 0.003), high *C_i_*-induced suppression of *C_i_*-independent stomatal opening was not observed for *ca1ca4* and Δ*g_s_* was similar across all *Ci* values >75 µmol mol*^-1^* (P < 0.05, for all Tukey’s comparisons see Table S3).

### Testing the relationship between PQ redox and *C*_i_-independent red-light responses

To address the hypothesised relationship between plastoquinone redox and red light-induced stomatal movements, we analysed the relationship between 1 – q*_L_* (Q*_A_* redox) and the *C_i_*-independent component of the red-light response curves in Col-0 (Figure 5). Interestingly, a more linear relationship was observed between Q *_A_* redox and *g_s_* (Figure 5) compared to the relationship between *g_s_* and red-light intensity (Figure 1B), suggesting Q *_A_* redox could be a better predictor of *g_s_*. Multiple linear regression in combination with backwards stepwise elimination yielded a minimal model for prediction of *g_s_* with significant main effects of Q *_A_* redox, *C_i_* and their interaction, which was able to capture a significant proportion of the experimental variance (*R^2^* =0.8305, *F*(13, 364)=137.2, *P* < 0.001; see Table S3). The significant interaction between Q *_A_* redox state and *C_i_* value reflected the fact that at *C_i_* values below 375 µmol mol *^-1^* individual regression lines yielded similar slopes (Figure 5A, Table S3), while at *C_i_* values above 460 µmol mol*^-1^*, slopes of the regression lines were significantly decreased (Figure 5A, Table S4), due to the observed stomatal closure response elicited by high *C_i_* which counteracted the effect of *C_i_*-independent red-light opening responses (Figure 1B & 2B).

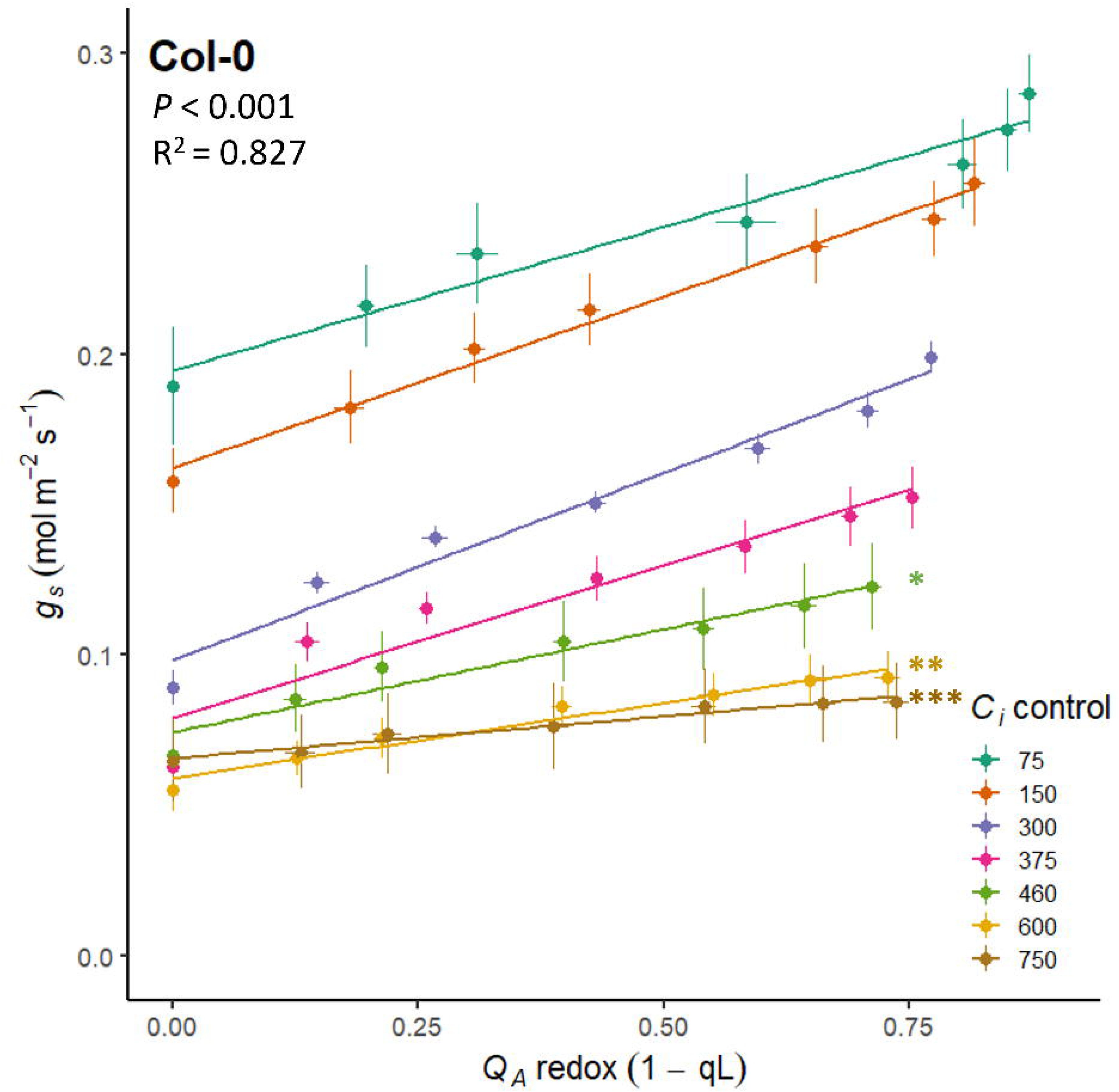

Intriguingly, the CO*_2_* hyposensitivity of *ca1ca4* caused the positive relationship between Q*_A_* redox and *g_s_* to be maintained across all measured *C_i_* values (Figure 6). As a result, multiple linear regression followed by backwards stepwise elimination provided a minimum model that did not include an interaction term, but only significant main effects of *C_i_* and Q*_A_* redox state on stomatal conductance (Figure 6, Table S5). This indicates that in the absence of high *C_i_*-induced stomatal closure, the modelled effects of Q*_A_* redox state and *C_i_* on *g_s_* become additive across the full range of *C_i_* concentrations.

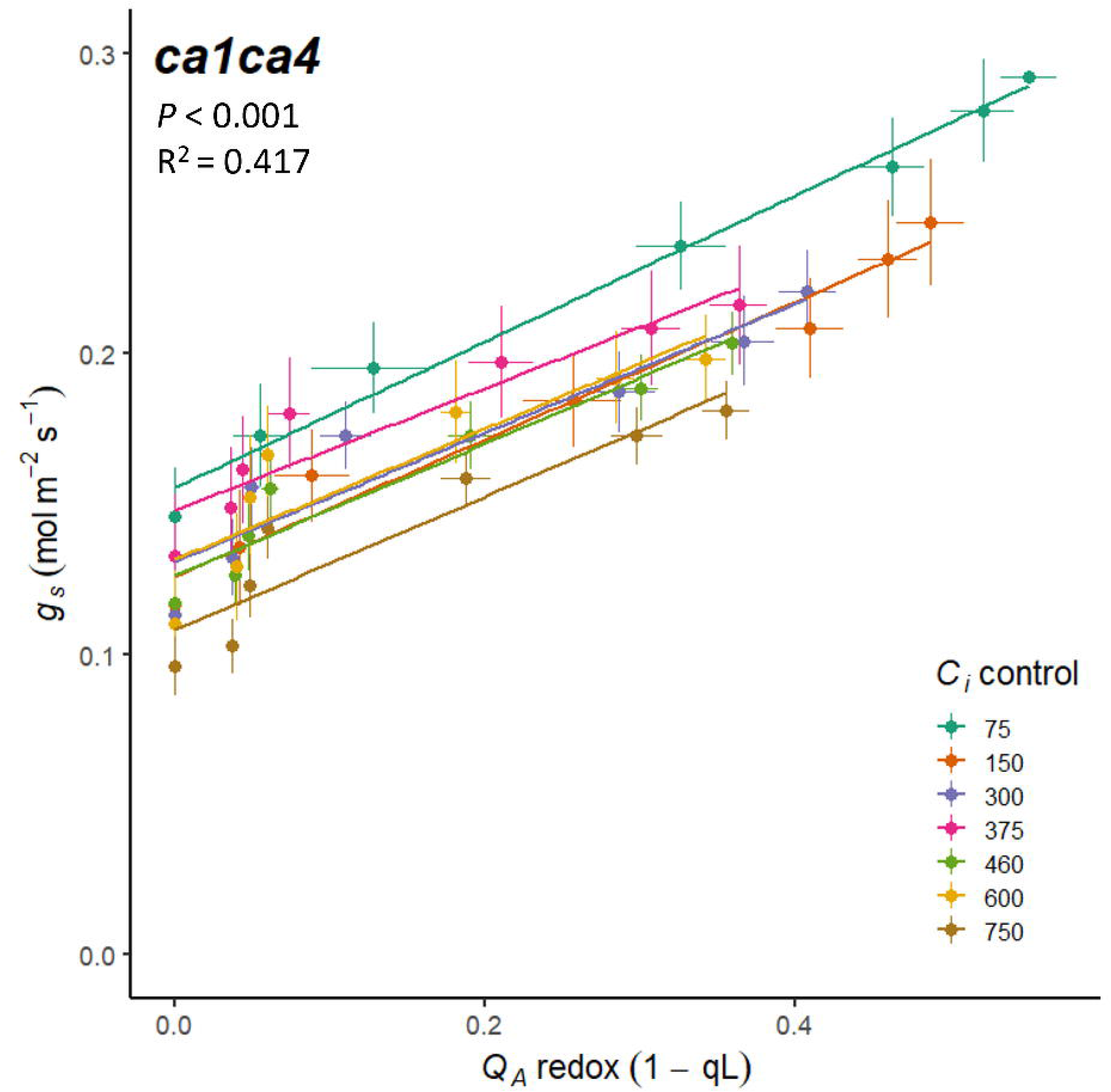

## Discussion

In this work, we quantitatively assessed the impact of *C_i_*-dependent and *C_i_*-independent mechanisms on the stomatal red-light response. By keeping *C_i_* constant across red light dose response curves at values spanning from 75 to 750 µmol mol *^-1^*, including concentrations within a ‘physiologically active range’, we were able to decouple *C_i_-*dependent from *C_i_*-independent mechanisms in coordinating stomatal behaviour. Subsequent analysis of these responses in the CO *_2_* hyposensitive mutant *ca1ca4* allowed further deconvolution of the involvement of these two red light-associated mechanisms.

### *C*_i_-dependent and -independent mechanisms contribute equally to stomatal red light responses

The control of *C_i_* over stomatal movements is well-known (Roelfsema *et al*., 2002, 2006) and it is increasingly clear that both *C_i_-*dependent and -independent mechanisms coordinate the stomatal red-light response (Messinger*et al.,* 2006; Lawson *et al.,* 2008; Wang & Song, 2008; Matrosova *et al.,* 2015). However, quantitative knowledge regarding the proportional contribution and extent of crosstalk between *C_i_*-dependent and -independent pathways has so far remained limited. Somewhat surprisingly, our results indicate that the relative magnitude of *C_i_-*dependent and -independent contributions to the stomatal red-light response is approximately equal (Figure 2B). The experimental design employed here allowed easy separation of *C_i_-*dependent and -independent components. While the response of *g_s_* to the controlled *C_i_* value during each red-light response curve confirmed the established role of *C_i_*-dependent stomatal movements (Roelfsema *et al*., 2002, 2006; Assman & Jegla, 2016; Engineer *et al.,* 2016), stomata also continued to respond to increasing red-light intensity when *C_i_* was held constant across a range of values (Figure 2B), clearly supporting a stomatal response to red light that is independent of *C_i_*, as previously observed (Messinger *et al.,* 2006; Lawson *et al.,* 2008; Wang & Song, 2008).

### Is there crosstalk between *C*_i_-dependent and -independent stomatal red-light responses?

Despite pronounced differences in the absolute values of *g_s_*, the overall shape and magnitude of the stomatal response (Figure 1B & 2B) remained similar between most *C_i_* values; suggesting the pathways which govern the *C_i_-*dependent and -independent responses are additive and at least partially independent of one another. The observed stomatal opening response to low *C_i_* in darkness, i.e., in the absence of a red-light signal (Figure 1B & 2A) provides further evidence that the *C_i_-*dependent mechanism is independent from alternative red-light-driven stomatal responses. The observed dark response is consistent with previous work, where although less pronounced and somewhat delayed, stomatal opening in response to low CO *_2_* was also observed under dark conditions (Sharkey & Raschke, 1981; Messinger *et al.,* 2006; Hiyama *et al.,* 2017; Hosotani *et al.,* 2021). Notably, this low-CO*_2_*-induced stomatal opening in the absence of light was abolished in plants lacking HT1 and CBC1/CBC2, suggesting HT1-CBC mediated suppression of S-type anion channels is required to facilitate stomatal opening in the dark (Hiyama *et al.,* 2017). Interestingly, Hiyama *et al*. (2017) also showed that while stomata do respond to low CO *_2_* in the dark, there was no further stomatal closure observed in response to high CO *_2_* (800 µmol mol*^-1^*). This is also consistent with the present study, as*g_s int_* was similar between all *C_i_* concentrations greater than 375 µmol mol *^-1^*, suggesting stomata are already maximally closed under ambient CO *_2_* in the dark. Notably, at *C_i_* values equal to those measured at high-light*C*(*_i_ =* 300 µmol mol*^-1^*) an intermediate*g_s int_* was observed, which suggests the presence of a threshold *C_i_* concentration, below which the opening response to low *C_i_* overrides other pathways of dark-induced closure. Interestingly, the *C_i_*-independent response estimated by Δ *g_s_* was not affected by the initial dark conductance. For example, for the stomatal red-light response determined at a *C_i_* value of 75 µmol mol*^-1^*, *g_s int_* was higher than the final *g_s max_* (at 800 µmol m*^-2^* s*^-1^*) observed for *C_i_* values above 300 µmol mol *^-1^*, but despite this, Δ*g_s_* remained similar (Figure 2A). As such, it could be postulated that the *C_i_-*dependent pathway functions as a ‘coarse’ baseline control of stomatal movements, whereas the *C_i_*-independent stomatal response to red light could be involved in further fine-tuning, to facilitate a more direct relationship with concurrent photosynthetic rates (Lawson *et al.,* 2008; Messinger *et al.,* 2006). In contrast to the independent operation of *C_i_*-dependent and -independent responses at low *C_i_*, the magnitude of Δ*g_s_* was significantly suppressed under high *C_i_* concentrations (Figure 2B). Thus, it seems that high *C_i_-*induced stomatal closure is dominant and suppresses *C_i_*-independent red-light-induced stomatal opening. The observed competition between both pathways is consistent with recent work where high CO*_2_*-mediated dephosphorylation of guard cell PM H *^+^*-ATPases was delayed and required a higher *C_i_*under red light compared to in darkness (Ando *et al.,*2022), indicating that red light-induced phosphorylation of PM H *^+^*-ATPases is a component of the*C_i_*-independent response (Ando *et al.,* 2022).

Interestingly, in contrast to the observations in Col-0, the CO *_2_* hyposensitive mutant *ca1ca4* displayed a similar *g_s int_* across all measured *C_i_* concentrations (Figure 4A). The response of *g_s int_* to low *C_i_* may be less visible in *ca1ca4* due to a diminished high *C_i_* response, and as such, could partially suppress dark-induced stomatal closure. The absence of a *C_i_*-dependent *g_s int_* response could also suggest that either carbonic anhydrase activity, or downstream HCO *_3_^-^* accumulation and sensing, or both, are required for the induction of low *C_i_*-induced stomatal opening in the absence of light. The impairment of low *C_i_*-induced stomatal opening is consistent with previous results, in which the response of *ca1ca4* to low CO*_2_* is delayed and much less pronounced (Hu *et al.,* 2010; Matrosova *et al.,* 2015). It has also been reported that the *C_i_*-independent red-light response of *ca1ca4* is stronger than the response to low CO*_2_* (Matrosova *et al.,* 2015). As such, it seems plausible that a concomitant light signal is required to evoke an opening response to low *C_i_* in these CO*_2_* hyposensitive plants. The inclusion of *ca1ca4* in the experiments reported in here enabled further isolation of *C_i_*-independent stomatal responses from the *C_i_-*dependent pathway. Interestingly, *ca1ca4* displayed a similar magnitude of Δ*g_s_* across all *C_i_* concentrations (Figure 4B). This sustained *C_i_-*independent opening response in *ca1ca4* can be attributed to the impairment of high CO *_2_*-induced stomatal closure (Hu*et al.,* 2010). Indeed, the high *C_i_*-induced suppression of stomatal opening observed for Col-0 (Figure 1B & 2B) was not present in *ca1ca4* (Figure 3B & 4B). Meanwhile, *C_i_*-independent mechanisms of stomatal opening remained unaffected in *ca1ca4* (Hu *et al.,* 2010; Matrosova *et al.,* 2015; Ando *et al.,* 2022); further supporting the idea that the red-light specific *C_i_*-independent stomatal opening response is at least partially independent of the mechanisms driven by *C_i_*.

### What signalling pathways coordinate the *C*_i_-dependent and -independent responses?

Overall, it seems that while the signalling pathways of the *C_i_-*dependent and -independent stomatal red-light responses may partially overlap, other components of these pathways are likely distinct. Components such as HT1, CBC1*/*2 and SLAC1 appear likely candidates for crosstalk between the two pathways, as mutants display perturbed responses to both red light and *C_i_* (Hiyama *et al.,* 2017; Laanemets *et al*., 2013; Matrosova *et al.,* 2015). Furthermore, experiments by Ando and Kinoshita (2018) showed that while reductions in *C_i_*promoted PM H*^+^* ATPase activation in red-light illuminated samples, low *C_i_* in darkness failed to induce phosphorylation required for activation, demonstrating the need for both stimuli. Interestingly, in isolated epidermal peels the combination of low *C_i_* and red-light illumination also failed to induce stomatal opening, suggesting a role for a much-debated mesophyll-derived signal (see review by Lawson *et al*., 2014).

In contrast to the shared signalling components above, a number of molecular players appear to have a distinct role in the *C_I_*-dependent pathway. For example, several CALCIUM DEPENDENT PROTEIN KINASES (CPKs) appear to exclusively respond to CO *_2_*; the *cpk3/5/6/11/23* quintuple mutant showed impaired stomatal responses to high and low CO *_2_*, but stomatal opening in response to red light was unaffected (Schulze *et al.,* 2021). Likewise, in the double carbonic anhydrase *ca1ca4* mutant the stomatal response to CO *_2,_* but not red light, was diminished (Ando *et al.,* 2022; Matrosova *et al.,* 2015). Interestingly, PM H*^+^*ATPase phosphorylation and stomatal opening under red light were both increased in *ca1ca4*, compared to wild type (Ando *et al.,* 2022), which may imply that plants could compensate for the deficiency in the *C_i_*-dependent pathway by enhancing the *C_i_*-independent pathway to maintain effective carbon gain. This raises the question as to what signals are involved in sensing and communicating the red-light stimulus in the *C_i_*-independent pathway. By isolating the*C_i_*-independent responses across a range of *C_i_* values, we could establish strong linear correlations between Q*_A_* redox and *g_s_* (Figure 5, Table S4), with similar regression slopes across ambient and sub-ambient *C_i_* values (Figure 5). These results provide support for the hypothesis that PQ redox state could be an early signal in the *C_i_*-independent pathway (Busch, 2014). The maintenance of this positive linear correlation across all measured *C_i_* values in *ca1ca4* (Figure 6, Table S5) provides further support for the inclusion of PQ redox as a predictor of red light-induced stomatal movements. The involvement of PQ redox state is also consistent with the abolishment of red-light-induced opening by DCMU, which blocks electron transfer upstream of PQ, maintaining an oxidised PQ pool (Sharkey & Raschke, 1981; Olsen *et al.,* 2002; Messinger *et al.,* 2006; Wang *et al.,* 2011; Ando & Kinoshita, 2018) and observed stomatal responses in *PsbS* overexpression and silencing *N. tabacum* lines (Głowacka *et al.,* 2018) which perturb PQ redox state via altered excitation pressure at photosystem II. Though it seems PQ redox could provide an early cue in the stomatal red-light response, all empirical evidence accumulated thus far is largely correlative, making establishment of cause and effect difficult. In addition, it remains unclear whether a putative chloroplast-derived signal would originate from the mesophyll (Roelfsema *et al.,* 2002; Lawson *et al.,* 2014) requiring further transduction to the GCs or within the GCs themselves (Zeiger, 1983; Olsen *et al.,* 2002). While the fluorescence measurements used here to estimate Q *_A_* redox state (Figure 1F & 3F) predominantly originate from the mesophyll, a mechanism based on signals derived from GC chloroplasts in coordinating stomatal movements appears more parsimonious and cannot be ruled out.

### Conclusions

Measurements of *g_s_* as a function of a matrix of *C_i_* values and red-light intensities allowed easy isolation of *C_i_*-dependent and -independent mechanisms in stomatal responses to red light and for the first time provided a quantitative assessment of the relative magnitude of each pathway. Results indicate that both pathways contribute equally to red-light stomatal responses, but high *C_i_* seems to override *C_i_*-independent opening responses. Abolishment of high *C_i_*-induced stomatal closure in *ca1ca4* removed the significant interaction between both mechanisms, giving rise to a model with two distinct additive *C_i_*-dependent and -independent mechanisms jointly coordinating stomatal opening in response to red light. While our data are consistent with the involvement of plastoquinone redox state in coordinating stomatal responses to red light, they highlight areas for further research to fully characterise additional signalling components involved in the regulation of the *C_i_*-independent stomatal red-light response.

## Acknowledgements

The authors thank Dr Rich Vath for general support with gas exchange measurements and Dr Jessica Royles for technical support. For the purpose of open access, the authors have applied a Creative Commons Attribution (CC BY) licence to any Author Accepted Manuscript version arising from this submission.

## Supplemental Materials

**Table S1** Genotyping primers used in this study

**Table S2** Dunn’s pairwise comparison of the initial dark conductance (*g_s int_*) and the total stomatal response (Δ*g_s_*) in *A. thaliana* Col-0.

**Table S3** Dunn’s pairwise comparison of the initial dark conductance (*g_s int_*) and the total stomatal response (Δ*g_s_*) in the CO*_2_* hyposensitive mutant *ca1ca4*.

**Table S4** Multiple linear regression analysis of the relationship between *g_s_* and Q*_A_* redox in Col-0

**Table S5** Multiple linear regression analysis of the relationship between *g_s_* and Q*_A_* redox in *ca1ca4*

**Figure S1.** Determining the response of *Ci* at a range of physiologically relevant red-light intensities.

## Author contributions

GT and JK designed the study. GT carried out the experiments with support from JW. GT and JK performed the data analyses. All authors helped to interpret the results. GT and JK wrote the paper with input from JW.

## Conflict of interest

The authors have no conflicts of interest to declare.

## Funding statement

GT was supported by the Alexander James Keith Studentship and the doctoral training programme of the School of Biological Sciences at the University of Cambridge. JK acknowledges new lecturer support from the Gatsby foundation.

## Data availability

All data supporting the findings of this study are available within the paper and within its supplementary materials.

## Notes

### Competing Interest Statement

The authors have declared no competing interest.

